# m^6^A landscape is more pervasive when *Trypanosoma brucei* exits the cell cycle

**DOI:** 10.1101/2024.02.23.581781

**Authors:** Lúcia Serra, Sara Silva Pereira, Idálio J. Viegas, Henrique Machado, Lara López-Escobar, Luisa M Figueiredo

**Affiliations:** Instituto de Medicina Molecular, Faculdade de Medicina, Universidade de Lisboa; Católica Biomedical Research Centre, Católica Medical School, Universidade Católica Portuguesa; Institute of Inflammation and Ageing, University of Birmingham

**Keywords:** m^6^A, immunoprecipitation, parasite, differentiation

## Abstract

N6-methyladenosine (m^6^A) is a mRNA modification with important roles in gene expression. In African trypanosomes, this post-transcriptional modification is detected in hundreds of transcripts and it affects the stability of the variant surface glycoprotein (VSG) transcript in the proliferating blood stream form. However, how m6A landscape varies across the life cycle remains poorly defined. Using full-length, non-fragmented RNA, we immunoprecipitated and sequenced m^6^A-modified transcripts across three life cycle stages of *Trypanosoma brucei* – slender (proliferative), stumpy (quiescent), and procyclic forms (proliferative). We found that 1037 transcripts are methylated in at least one of these three life cycle stages. While 21% of methylated transcripts are common in the three stages of the life cycle, globally each stage has a distinct methylome. Interestingly, 47% of methylated transcripts are detected in the quiescent stumpy form only, suggesting a critical role for m^6^A when parasites exit the cell cycle and prepare for transmission by the Tsetse fly. In this stage, we found that a significant proportion of methylated transcripts encodes for proteins involved in RNA metabolism, which is consistent with their reduced transcription and translation. Moreover, we found that not all major surface proteins are regulated by m^6^A, as procyclins are not methylated, and that, within the VSG repertoire, not all VSG transcripts are demethylated upon parasite differentiation to procyclic form. This study reveals that the m^6^A regulatory landscape is specific to each life cycle stage, becoming more pervasive as *T. brucei* exits the cell cycle.

**Summary:** African trypanosome parasites adapt to mammalian and insect hosts by adjusting gene expression, morphology, and metabolism. In this study, we focus on how N6-methyladenosine (m^6^A), a post-transcriptional modification, affects the parasite’s transcriptome throughout its differentiation from the mammalian host to the fly. We found that methylation is differentially regulated as the life cycle progresses, being particularly prevalent in the non-proliferative stumpy form, as more methylated transcripts are found at this insect-infective stage than in slender and procyclic forms. We further show that the not all parasite surface proteins are regulated by m^6^A and that the previously identified link between m^6^A methylation and the expression level of the major surface protein of bloodstream forms applies to the active variant surface glycoprotein, but not always to silent genes, suggesting two distinct regulatory mechanisms of (de)methylation.

## Introduction

*Trypanosoma brucei* is a unicellular parasite that causes sleeping sickness in humans and nagana in livestock [1]. The parasite’s life cycle alternates between a mammalian host and the tsetse fly, relying on a complex set of metabolic, morphological and gene expression adaptations to ensure transmission and survival in drastically different environments [2]. In mammals, *T. brucei* survives in the bloodstream as proliferative slender forms, which express a surface coat of variant surface glycoproteins (VSGs). Although the *T. brucei* genome encodes over 2000 *VSG* genes, only one *VSG* gene is expressed at any given time from the corresponding bloodstream expression site (BES) [3]. Selection and expression of a new *VSG* gene occurs after homologous recombination or transcriptional activation of another BES, allowing the parasite to regularly switch its VSG coat and thus escape the host immune system by antigenic variation [4].

When slender forms reach a critical cell density in the bloodstream, they differentiate into non-replicative stumpy forms through a quorum-sensing mechanism triggered by oligopeptides, generally known as the stumpy induction factor (SIF). Such peptides are generated by parasite-released peptidases in a cell density-dependent manner and activate a signaling pathway that leads to gene expression remodeling, including reduction of transcription and translation [5–8]. Uptake of bloodstream parasites by the tsetse fly triggers further differentiation into procyclic forms, which lead to broad transcriptomic and morphological changes, including the replacement of the VSG by a procyclin surface coat [9].

RNA modifications are important regulators of gene expression [10]. The most prevalent modified nucleotide in eukaryotic mRNA is m^6^A (or N6-methyladenosine), which in mammalian transcriptomes can be found mostly enriched in the 3′ untranslated region (UTR) and near the stop codon [11]. The function of m^6^A varies between organisms, including parasites, where it might be important for life cycle progression. In *Toxoplasma gondii*, m^6^A methylation is prevalent in asexual life cycle stages [12] and is involved in proper mRNA 3’ end processing, likely affecting developmental gene regulation [13]. In *Plasmodium falciparum*, m^6^A is highly developmentally regulated, mediating translational repression of transcripts involved in regulation of gene expression in blood stages [14]. In *T. brucei*, Liu *et al*. found that m^6^A modifications were present in mRNA transcripts in slender and procyclic forms. Their study showed how both life cycle stages differed in terms of methylated transcripts and identified in which region of the transcript (coding or untranslated) m^6^A peaks were located [15]. More recently, our lab identified m^6^A can also be found in the poly(A) tail, where it stabilizes VSG transcripts in slender forms and whose removal preceded VSG downregulation during parasite differentiation to procyclic forms [16]. However, whether m^6^A methylation is important for gene expression regulation in other stages of the *T. brucei* life cycle remains unknown.

In this work, we mapped methylated transcripts in three stages of the parasite life cycle. By using full length, non-fragmented RNA as input, we mapped any transcript that harbored m^6^A, regardless of its location within the transcript. In contrast to mammalian cells, in which distribution of m^6^A methylation appears to be similar between tissues [17, 18], in *T. brucei*, the m^6^A landscape changes across life cycle stages. We show that the stumpy form transcriptome has a larger proportion of methylated transcripts compared to slender and procyclic forms. Finally, we also show that while procyclin is not methylated, VSG transcripts are differentially m^6^A-regulated as not all VSG transcripts lose their m^6^A methyl group in similar temporal patterns.

## Materials and Methods

### Cell culture and cell-lines

*Trypanosoma brucei* bloodstream parasites, including slender and stumpy forms (*T. brucei* EATRO 1125 AnTat 1.1E 90–13 GPF::PAD1^3’UTR^, a transgenic cell line in which GFP is coupled to PAD1 3′ untranslated region (UTR) were cultured in HMI-11, containing 10% Fetal Bovine Serum (10270106, Gibco) and 0.5% Penicillin-Streptomycin (15070063, Gibco) at 37°C in 5% CO2.

Differentiation of slender into stumpy forms was performed by following two different protocols in parallel. One protocol induced differentiation from slender forms, at starting cell density of 5×10^5^ parasites/ml, by adding pCPT-cAMP (8-(4-chlorophenylthio)adenosine 3’:5’-cyclic monophosphate, C3912, Merck Life Sciences) in the HMI-11 culture to a final concentration of 10µM. Parasites were left to grow for 48h at 37°C in 5% CO2. The second protocol induced differentiation by growing slender forms, at starting cell density of 2×10^5^ parasites/ml, in HMI-11 containing 10% Fetal Bovine Serum, 1.1% methylcellulose (Methocel A4M, 94378, Merck Life Sciences), and 0.1% gentamicin (15750060, Gibco) at 37°C in 5% CO2 for 48h.

To differentiate stumpy into procyclic forms, stumpy forms were collected from culture 48h after pCPT-cAMP addition, spun down, and resuspended in SDM-79 [19] (RR110008P1, ThermoFisher) supplemented with 10mM glycerol (G5516, Sigma-Aldrich), 6 mM cis-aconitate[25] (A3412, Sigma-Aldrich) and 0.5% Penicillin-Streptomycin at 2×10^6^ parasites/ml. Procyclic formscwere then incubated at 27°C for 5 days [20].

### Microscopy

Small aliquots of slender, stumpy, or procyclic forms were collected from cultures and fixed for 5 min at room temperature with 2% of formaldehyde (F8775, Sigma-Aldrich). To stain the nuclei, 1µl of Hoechst solution (1ug/ml) was added to the fixed cells and immediately washed twice with 500 µL of 1X PBS. Parasites were then pelleted by centrifugation for 5 min at 769g and resuspended in 50 µL of 1X PBS. For each population, 50-100 µL were deposited on a SuperFrost™ Plus slide (Fisher Scientific) and allowed to adhere for at least 4 hours in a humid environment (to prevent desiccation). Afterwards, the supernatant was removed by decantation and 5µl of Fluoromount-G™ Mounting Medium (00-4958-02, Invitrogen) were added to the sample.

Imaging of the slender, stumpy, and procyclic form parasites was performed on a confocal point-scanning microscope with Airyscan Zeiss LSM 880 using a 63x objective lens (Plan-Apochromat, NA 1.40, oil immersion, Zeiss). Laser stacks 405 (405 nm) and 488 (488 nm) were used for visualizing kinetoplasts and nuclei (Hoechst) and GFP (Green Fluorescence Protein) respectively. Acquired images were then revised with ImageJ.

### Flow cytometry

Single-cell suspensions containing parasites from each slender and stumpy cell culture were prepared for analysis of PAD1 expression, cell cycle distribution and cell viability by flow cytometry. Viability analysis was performed by staining parasites in culture medium with 0.01 mg/ml of propidium iodide (P4864, Merck Life Sciences). For cell cycle and PAD1 expression analysis, 0.2–1×10^6^ cells were fixed with ice-cold ethanol and stained with 0.01 mg/ml of propidium iodide (P4864, Sigma-Aldrich), as previously described [21]. Samples were passed through a 40-µm pore size nylon cell strainer (352340, Corning) and then analyzed on a BD LSRFortessa flow cytometer with FACSDiva 6.2 Software. All data were analyzed using FlowJo software version 10.0.7r2. Statistical analysis was performed using GraphPad Prism (version 8.4.3). Statistical differences were assessed using two-sided unpaired or multiple t-tests. *p*-values lower than 0.05 were considered statistically significant.

### RNA isolation and handling

Per condition, at least 200 million parasites were collected. Slender and stumpy form parasites growing in methylcellulose were washed 5 times in warm 1X trypanosome dilution buffer (5 mM KCl, 80 mM NaCl, 1 mM MgSO4, 20 mM Na2HPO4, 2 mM NaH2PO4, 20 mM glucose, pH 7.4) and centrifuged at 2200g for 10 min at 37°C to dilute the methylcellulose. Subsequently, all slender, stumpy and procyclic cultures were lysed in 1 mL of TRIzol (15596026, Invitrogen). RNA was isolated according to the instructions of the manufacturer. RNA was treated with DNAse I (M0303, NEB) (1 U per 2.5 µg of RNA) for 1 h at 37°C. Reaction was inactivated by adding 5 mM EDTA (AM9260G, Invitrogen) and heating to 75°C for 20 min.

### m^6^A immunoprecipitation and sequencing (m^6^A-IP)

m^6^A-IP sequencing was performed on 20µg of DNAse-treated total RNA extracted from (i) slender forms in HMI-11; (ii) slender forms in HMI-11 supplemented with 1.1% methylcellulose; (iii) stumpy forms derived from differentiation *in vitro* with pCPT-cAMP addition; (iv) stumpy forms derived from differentiation in HMI-11 supplemented with 1.1% methylcellulose; (v) procyclic forms grown in SDM-79 supplemented with 10mM glycerol.

Briefly, protein A/G magnetic beads (50 µL per sample, ThermoFisher, 88802) were washed in IP buffer (150 mM NaCl, 10 mM Tris-HCl, pH 7.5, 0.1% NP-40) three times. The beads were incubated with anti-m^6^A antibody (5 µl, Cell Signalling D9D9W) in 500 µL of IP buffer at 4°C during 30 min with gentle agitation. After incubation, beads were washed three times with IP buffer. RNA samples were denatured at 75°C for 5 min and cooled on ice for 2-3 min. RNA was mixed with the conjugated beads and incubated 30 min at 4°C with gentle agitation. After incubation, beads were washed twice in IP buffer, twice in low salt buffer (50 mM NaCl, 10 mM Tris-HCl, pH 7.5, 0.1% NP-40), twice in high salt buffer (500 mM NaCl, 10 mM Tris-HCl, pH 7.5, 0.1% NP-40) and twice in IP buffer. After the last wash, beads were resuspended in 400 µL of RLT buffer (Qiagen, cat. 79216s) and RNA was purified with RNeasy mini kit (Qiagen, cat. 50974104). RNA concentration and integrity were checked by fluorometry (Qubit RNA HS, Thermo Fisher Scientific) and parallel capillary electrophoresis (Fragment Analyzer, BioLabTech), respectively. cDNA was prepared and amplified using the SmartSeq2 protocol (Illumina, USA) and sequencing libraries were prepared with Pico Nextera kit (Illumina), as per manufacturer instructions. cDNA was sequenced as 75bp single-end reads on the NextSeq 550 platform (Illumina).

### Read processing and transcript identification

Reads were trimmed and mapped to the *T. brucei* EATRO 1125 genome (obtained from www.tritrypdb.org, release 65) using HISAT2 2.1.0 [22] under default settings. Alignment files were parsed through SAMtools (v1.17) [23], and transcripts were assembled and their count estimated using StringTie 1.3.6 [24], following the developers’ protocol [25]. To avoid missing VSGs, multi-mapping was allowed. Transcripts with less than 10 read counts throughout samples were filtered using edgeR [26] and enrichment levels were estimated with limma-voom [27], following a differential expression analysis pipeline. Transcripts were considered methylated when enriched in the m^6^A-IP samples, compared to the input. Enrichment was considered significant if the log2 fold change of m^6^A-IP / input (log2FC) was higher than 1 and the adjusted *p*-value was smaller than 0.05. To counteract transcript redundancy introduced by multi-mapping, enriched transcripts with more than 95% nucleotide identity to longer transcripts were removed. Gene set enrichment analysis was conducted in the GSEA software [27, 28] from the Broad Institute. Remaining statistical and clustering analyses were conducted in R.

## Results

### Generation and characterization of three parasite life cycle stages

In this study, we aimed to identify m^6^A-methylated transcripts in three different stages of *T. brucei’s* life cycle, regardless of the position of m^6^A within the transcript. Slender, stumpy and procyclic forms can be readily obtained in large amounts *in vitro*. Given the variety of experimental protocols available to obtain stumpy forms *in vitro*, we decided to use two protocols in parallel (Fig. 1A). In the first method, we cultured *T. brucei* EATRO 1125 slender forms for two days in the presence of pCPT-cAMP, a cyclic adenosine monophosphate (cAMP) analog that induces slender differentiation to stumpy forms (ST-CPT) [29]. The second method consisted in culturing slender forms for two days in 1.1% of methylcellulose (ST-MC). Methylcellulose increases the viscosity of the medium, mimicking the interaction of the parasite with the environment [30]. This method normally leads to a lower number of aberrant cell division phenotypes [31]. To obtain procyclic forms, we incubated stumpy forms obtained from the pCPT-cAMP differentiation protocol, in SDM-79 medium supplemented with glycerol and cis-aconitate at 27°C for five days (PCF) [20]. Slender forms were the starting population of parasites, and they were kept below a cell density of 5×10^5^ parasites/mL in either standard culture medium (SL) or medium supplemented with 1.1% methylcellulose (SL-MC).

**Fig. 1.**
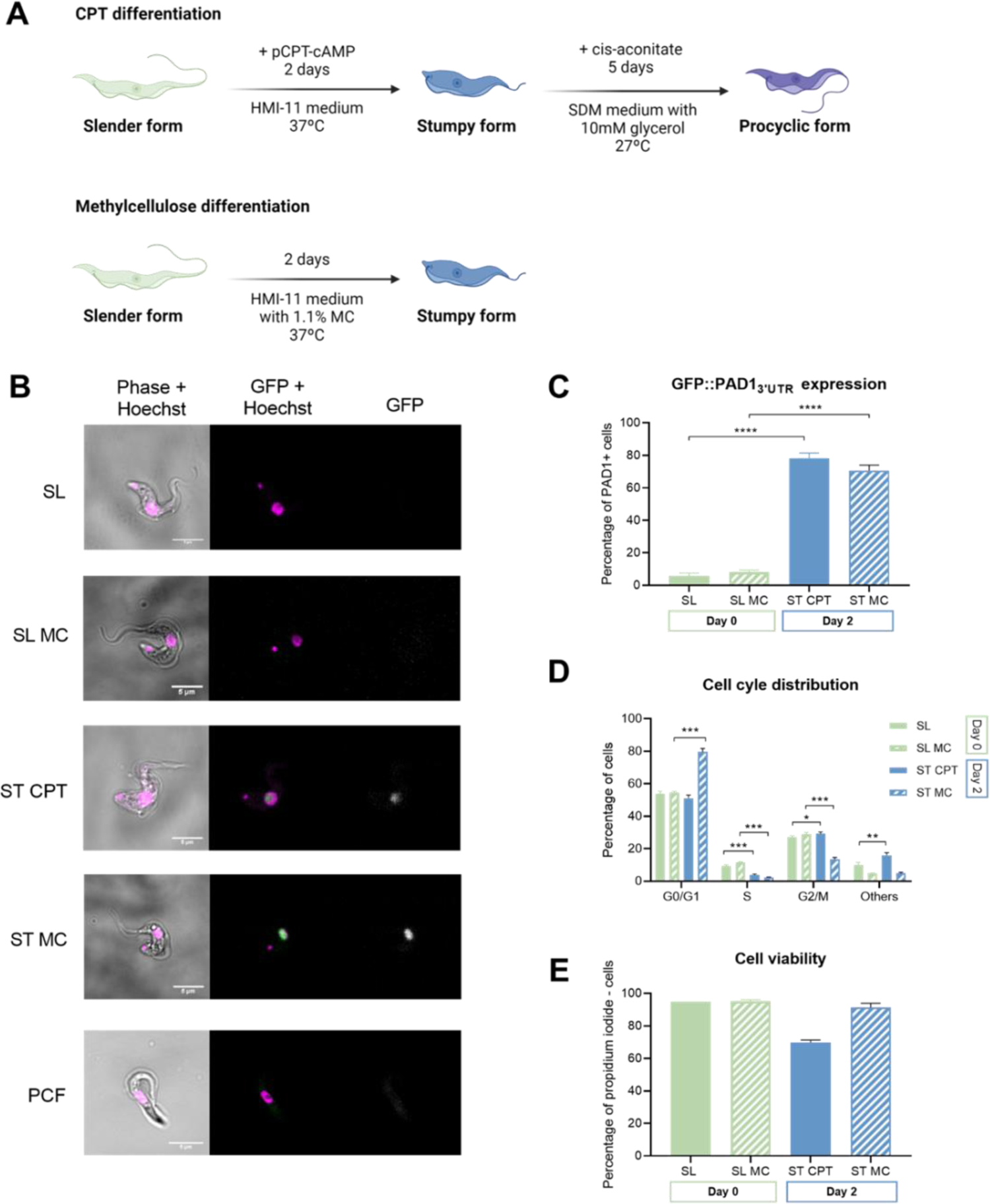
Profile of stumpy forms produced upon incubation with methylcellulose or cAMP analog. **A**, Experimental outline of two protocols performed in parallel for the *in vitro* differentiation of EATRO 1125 AnTat 1.1E 90–13 GPF::PAD1^3’UTR^cslender to stumpy parasites, using medium supplemented with either pCPT-cAMP (CPT) or methylcellulose (MC). Differentiation to procyclic forms was performed from pCPT-cAMP stumpy form parasites. Created with BioRender om. **B**, Representative images fromcslender (SL), stumpy (ST) and procyclic (PCF) form parasites collected from *in vitro* differentiation protocols (see panel A), n=4. Nuclei were stained by Hoechst. GFP signal corresponds to PAD1 marker expression. Scale bars, 5 µm. **C**, PAD1 expression analysis of Day 0 slender (SL and SL-MC) and Day 2 stumpy (ST-CPT and ST-MC) form parasites. Unpaired t-tests for Day 0 SL vs Day 2 ST-CPT; and for Day 0 SL-MC vs Day 2 ST-MC, *p*-value=0.0001). Error bars represent mean ± S.D, n = 4. **** *p*-value < 0.0001. **D**, Cell-cycle analysis of Day 0 slender (SL and SL-MC) and Day 2 stumpy (ST-CPT and ST-MC) form parasites. Unpaired t-test for Day 0 SL vs Day 2 ST-CPT at G0/G1, *p*-value=0.079; Unpaired t-test for Day 0 SL-MC vs Day 2 ST-MC at G0/G1, *p*-value <0.001) Unpaired t-test for Day 0 SL vs Day 2 ST-CPT atcOthers, *p*-value= 0.005; Unpaired t-test for Day 0 SL-MC vs Day 2 ST-MC at Others,c*p*-value=0.918). Error bars represent mean ± S.D., n = 4. * *p*-value < 0.05, ** *p*-value < 0.01, *** *p*-value< 0.001 **E**, Cell viability analysis assayed by flow cytometry of propidium iodide-stained Day 0 slender forms (SL and SL-MC) and Day 2 stumpy forms (ST-CPT and ST-MC). Error bars represent mean ± S.D., n = 4.

To confirm that the differentiation protocols of stumpy and procyclic forms were successful, we used four assays: (i) morphological assessment by microscopy, (ii) quantification of GFP::PAD1 reporter expression by flow cytometry, (iii) cell cycle profile, and (iv) cell viability. The majority of cells within each population showed expected morphology: elongated and thin for slender forms; short and stocky for stumpy forms; and pointy for procyclic forms, with the flagellum starting from the mid-body (Fig. 1B).

Expression of “proteins-associated with differentiation” (PAD), including PAD1, can be used as a marker of parasites that have committed to differentiation to stumpy forms [6, 32]. Given that stage specific PAD1 expression is dependent on the 3’UTR [32], we used a stumpy form reporter cell line in which the GFP gene is under the control of the PAD1 3’ UTR (EATRO 1125 AnTat 1.1E 90–13 GPF::PAD1) (J. Sunter, A. Schwede, and M. Carrington, personal communication; [33]). Both protocols used to produce stumpy forms led to an increase in the mean PAD1 expression (Fig. 1C, mean ± SD 78% ± 3% for ST-CPT, and 71% ± 4% for ST-MC Cell cycle analysis revealed that, in methylcellulose, a larger proportion of the parasite population was in G0/G1 than with pCPT-cAMP supplementation (Fig. 1D, 80% ± 2% for ST-MC; 51% ± 2% for ST-CPT). Parasites growing in medium supplemented with pCPT-cAMP showed more cell division abnormalities than in methylcellulose conditions, as seen by the mean increase of parasites in the category ‘Others’ (>4N) (Fig. 1D for ST-CPT: 16% ± 1.5; for ST-MC: 10% ± 2). Consistently, the mean cell viability was lower when parasites were differentiated with pCPT-cAMP than in methylcellulose-supplemented medium (Fig. 1E, ST-CPT: 70% ± 2; for ST-MC 91% ± 3).

These results show that the two protocols allow the development of stumpy forms with expected morphological and molecular characteristics, even though the use of a cAMP analog may cause more cell death at population level.

### Identification of m^6^A-methylated transcripts in three life cycle stages

RNA was extracted from slender forms (SL and SL-MC), stumpy forms (ST-CPT or ST-MC) and procyclic forms (PCF). Methylated full length transcripts were immunoprecipitated. Without fragmentation, m^6^A-enriched samples and corresponding input samples (i.e. pre-immunoprecipitation) were amplified by SMART-seq2 and sequenced. Reads were mapped to the *T. brucei* EATRO1125 genome, transcript counts were estimated and transcripts with low read counts were filtered (Fig. 2A). Multidimensional scaling showed a clear separation between transcriptomes of input and m^6^A-IP samples (Fig. 2B), as evidenced by the first principal component, which explains for the majority of the variance observed. This indicates that m^6^A immunoprecipitation was consistent across samples. Samples further clustered by life cycle stage, and to a lesser degree by differentiation protocol.

**Fig. 2.**
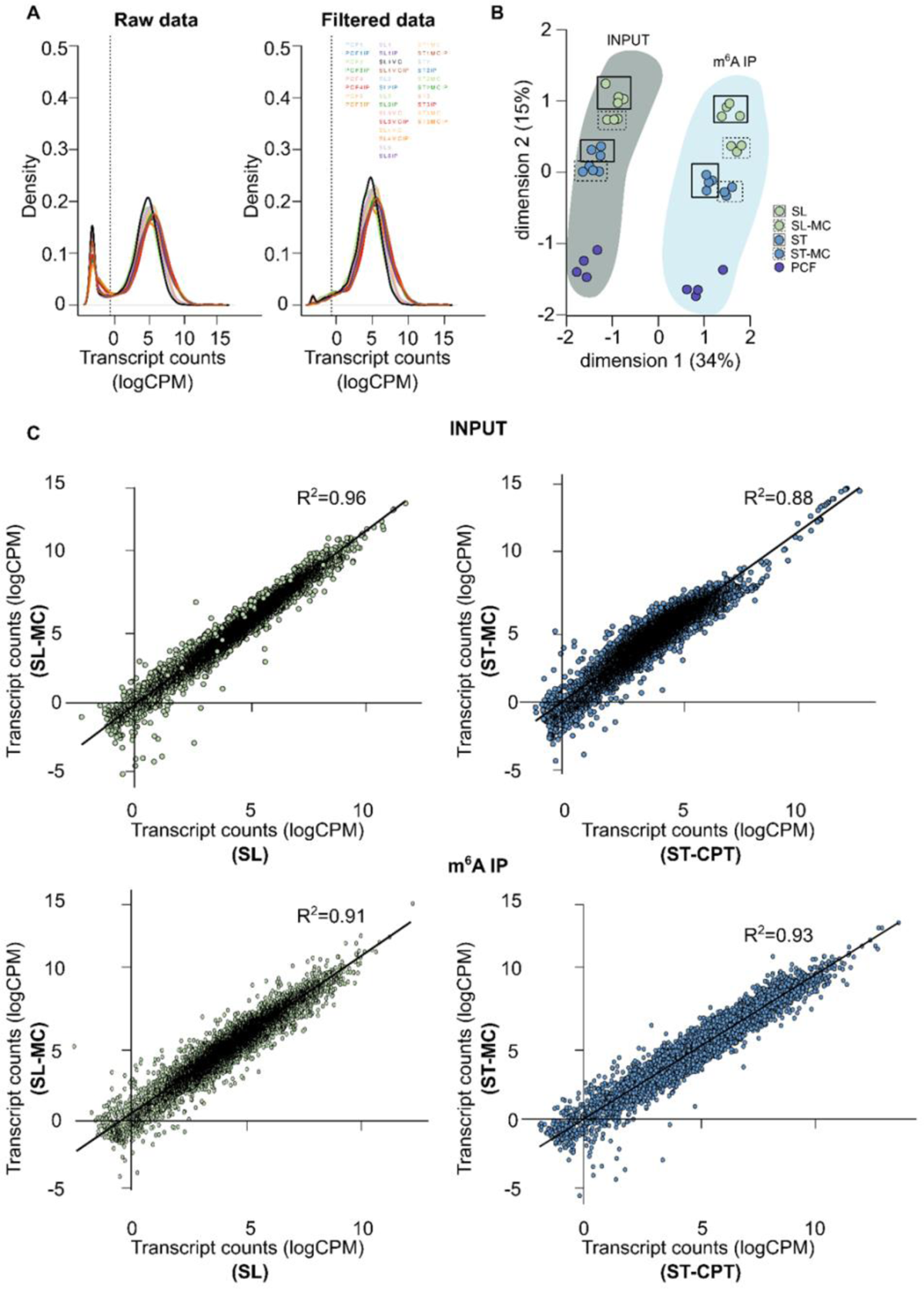
Transcriptome relationships before and after m^6^A enrichment. **A**, Transcript count density expressed as log-counts per million reads mapped (logCPM) per sample before and after filtering transcripts with less than 10 read counts across samples. **B**, Multidimensional scaling plot showing the distances between gene expression profiles of each experimental group: input (dark shade) and m^6^A-IP (light shade) samples separate across dimension 1, whilst samples from different parasite forms spread across dimension 2. Full boxes indicate slender form parasites grown in HMI-11 medium, whereas dashed boxes highlight samples grown in methylcellulose medium. Samples are color-coded according to key. **C**, Correlation between input (top) or m^6^A-IP (bottom) transcriptomes of slender (left, green) or stumpy (right, blue) form parasites grown in either HMI-11 or methylcellulose. Lines-of-best-fit are shown in black. R^2 values were estimated by Pearson’s correlation test.

To assess the impact of the medium in the transcriptomes (input samples) and methylomes (m^6^A-IP-samples), we compared the transcriptomes of slender form parasites grown in standard medium without methylcellulose (SL) with those grown in methylcellulose (SL-MC). We observed a high positive correlation between conditions, both from input and m^6^A-IP samples (Pearson’s R^2^ ranging between 0.96 and 0.91). The same tendency was observed when we compared stumpy forms obtained by pCPT-cAMP-induction (ST-CPT) with those differentiated by density in methylcellulose-supplemented medium (ST-MC) (Pearson’s R^2^ ranging between 0.88 and 0.93) (Fig. 2C). Given the low cell viability of ST-CPT population (Fig. 1E), it was not surprising to find a lower correlation in this comparison (Pearson’s R^2^ 0.88). In fact, 153 genes were differentially methylated between stumpy forms grown in normal medium *vs.* in medium supplemented with methylcellulose (Supplementary File A, table A1). Taking these results into account, in subsequent analyses we proceeded only with slender and stumpy form samples grown in methylcellulose.

For each life cycle stage, in the pre-immunoprecipitation RNA samples we identified a total of 9248 (slender), 9292 (stumpy), and 9212 (procyclic) different transcripts (Supplementary File A, table A2, A3 and A4 respectively). Subsequently, we compared transcript abundances before and after m^6^A immunoprecipitation, which allowed us to identify the m^6^A-methylated transcripts (Fig 3, Supplementary File B, tables B1, B2, B3). In total, we identified 1037 m^6^A-methylated transcripts in *T. brucei* EATRO1125 (Fig. 3A). Stumpy form is the life cycle stage with the highest number of methylated transcripts (968, 93% of all methylated transcripts), followed by slender forms (507, 49% of all methylated transcripts) and procyclic forms (262, 25% of all methylated transcripts) (Fig. 3A). Transcripts from 215 genes (21%) are methylated throughout the three life cycle stages, 234 (23%) are methylated in slender and stumpy forms only and 552 (53%) are life cycle stage specific.

**Fig. 3.**
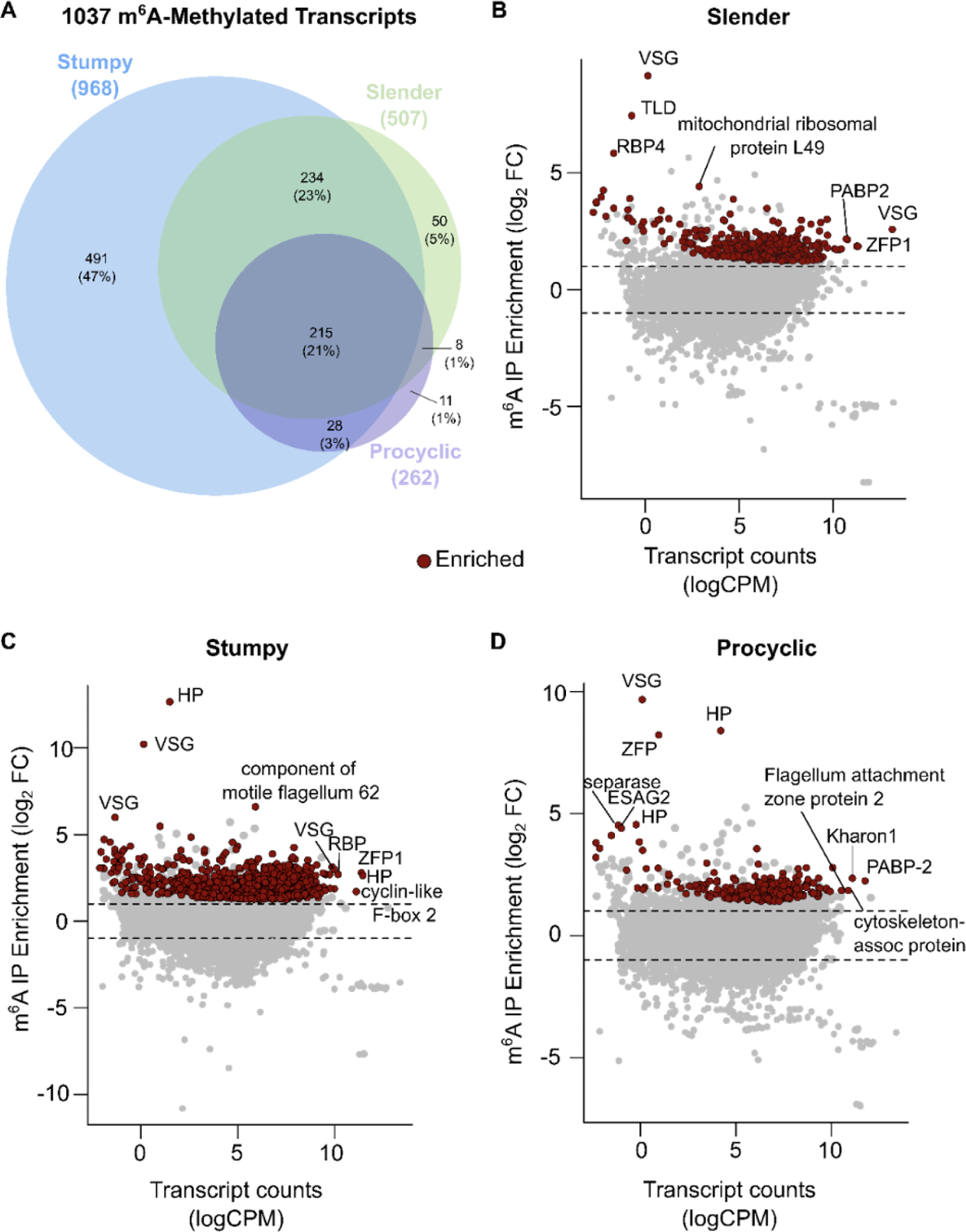
m^6^A landscape in slender, stumpy and procyclic forms. **A**, Venn diagram showing the number of m^6^A-enriched transcripts in slender, stumpy, and procyclic form parasites, and their intersections. **B**, MA plot showing transcripts differentially enriched before and after m^6^A-IP in slender form parasites. **C**, MA plot showing transcripts differentially enriched before and after m^6^A-IP in stumpy form parasites. **D**, MA plot showing transcripts differentially enriched before and after m^6^A-IP in procyclic form parasites. The y axis depicts m^6^A enrichment, shown as log2 fold change between m^6^A-IP and input samples. The x axis shows mean transcript counts, represented as log-counts per million reads mapped (logCPM). Significantly changed genes (log2FC >|1| and *p*-value < 0.05) are highlighted in red (enriched in m^6^A-IP).

As previously reported for strain Lister 427, VSG transcripts were amongst the most enriched transcripts in slender forms [16], together with a TLD (Tre2/Bub2/Cdc16 (TBC), lysin motif (LysM) domain-containing protein, RNA binding protein 4, and mitochondrial ribosomal protein L49. Within the most abundant transcripts, we found the active VSG, Zinc-finger protein 1, and polyadenylate-binding protein 2 (PABP2) also enriched in m^6^A (Fig. 3B). VSGs were also amongst the transcripts most enriched for m^6^A in stumpy forms, together with a gene encoding for a component of motile flagellum 62 (Fig. 3C). Consistent with previous studies, in this life cycle stage the active VSG was abundant, but not the most abundant transcript [8]. We also detected another RNA binding protein, a zinc-finger protein, a hypothetical protein, and a cyclin-like F-box 2 (CFB2) protein within the most abundant transcripts. Analysis of transcripts from procyclic forms showed again a silent VSG as the transcript most enriched in m^6^A, followed by a hypothetical protein, a zinc-finger protein, separase and one expression site-associated gene 2 (ESAG2) (Fig. 3D). Within the most abundant transcripts also enriched for m^6^A, we detected cytoskeleton-associated protein, Kharon 1, flagellum attachment zone protein 2, and again PABP-2. Procyclin, despite its high abundance, was not amongst the transcripts enriched for m^6^A.

Overall, we identified m^6^A-methylated transcripts in slender, stumpy and procyclic forms. Comparing with previous studies, we observed that the methylated transcriptome (methylome) is globally reproducible (Supplementary File B, tab B4). Here, the coverage is deeper, we potentially retrieved both poly(A) and internally methylated transcripts, and we directly compared three stages of a pleomorphic strain of *T. brucei*.

### m^6^A methylation in three stages of life cycle

In this study, we considered all methylated transcripts found in any of the three life cycle stages (log fold change>1 and adjusted *p*-value < 0.05, N=1001) as the full methylome. To analyze the dynamics of m^6^A methylation across the three life cycle stages and the functions of the methylated transcripts, we divided the methylated transcripts into 7 clusters based on their enrichment in each life cycle stage (Fig. 4A). Cluster 1 contains transcripts methylated in the three life cycle stages (N=215), thus constituting the core methylome; in cluster 2 (N=44) we grouped transcripts methylated only in slender forms; cluster 3 (N=470) only in stumpy forms; and cluster 4 (N=11) only in procyclic forms. Cluster 5 (N=225) contains transcripts methylated in slender and stumpy forms, but not procyclic forms; cluster 6 (N=8) comprises transcripts methylated in slender and procyclic forms, but not stumpy forms; finally, in cluster 7 (N=28), we pooled transcripts methylated in stumpy and procyclic forms, but not slender forms.

**Fig. 4.**
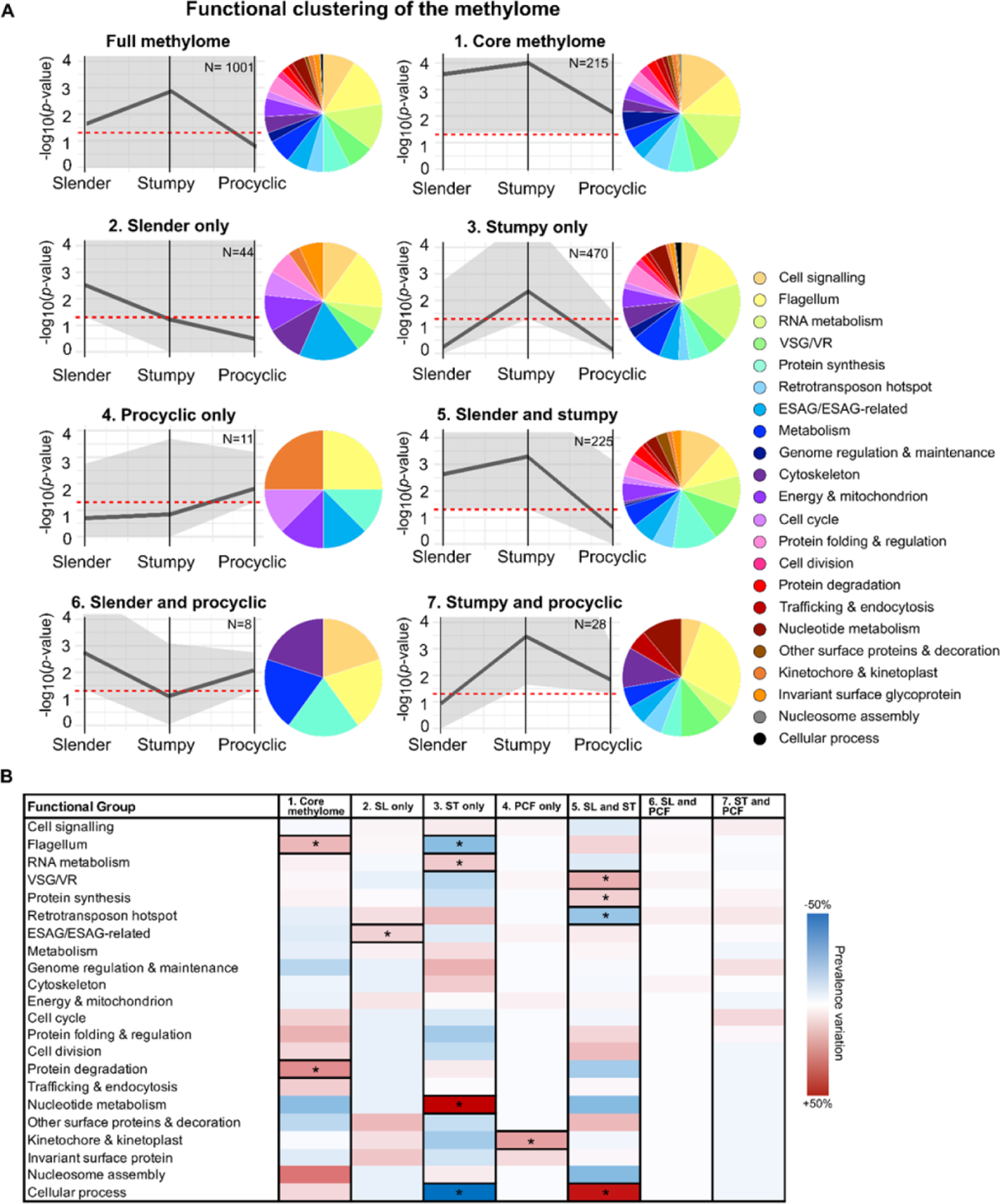
Clustering and functional characterization of methylated transcripts in three stages of *T. brucei* life cycle. **A.** From the pool of transcripts significantly enriched in at least one life cycle stage (full methylome, N=1001), we built 7 clusters: cluster 1, or core methylome, contains genes methylated across the three life cycle stages (N=215); clusters 2 (N=44), 3 (N=470), and 4 (N=11), contain genes methylated exclusively in slender, stumpy, or procyclic form parasites, respectively; cluster 5 (N=225) contains genes methylated in slender and stumpy form parasites, but not procyclic forms; cluster 6 (N=8) contains genes methylated in slender and procyclic form parasites, but not stumpy forms; cluster 7 (N=28) contains genes methylated in stumpy and procyclic forms, but not slender form parasites. Grey lines show mean m^6^A enrichment; grey shades show range of m^6^A enrichment values; red lines mark m^6^A enrichment cutoff (log2FC >1). Each transcript from each cluster was assigned one of 22 functional categories and color-coded according to key. **B.** Assessment of the size variation of each functional group across the 7 clusters. Thick-frame boxes indicate statistically significant changes, color-coded according to key.

To functionally characterize the transcripts present in each cluster, first, we removed the transcripts for which functional information was not available in VEuPathDB (i.e. hypothetical proteins), corresponding to 32% ± 3% (mean ± SD) of the total. Given that GO term annotation in *T. brucei* is limited, we assigned each transcripts to one of 22 manually curated functional groups (Fig. 4A, legend panel). The members of individual clusters and their functions can be found in Supplementary File C.

From this functional analysis, we conclude that m^6^A methylation occurs in transcripts encoding for proteins with a wide variety of functions and subcellular localizations. Notably several transcripts of a given function are differentially methylated and do not follow the same pattern of methylation across the three stages of the parasite’s life cycle. Changes in the pattern of m^6^A methylation do not appear to be restricted to a subset of functionally related genes in any life cycle stage but are found instead across all functional categories. Interestingly, among the methylated transcripts exclusively methylated in stumpy forms, one of the most represented functions is RNA metabolism, which could be important for the gene expression rewiring that takes place in this stage [7, 8].

Next, we assessed how transcripts from each functional group distribute across each cluster. For this, we compared the prevalence in each functional group throughout the clusters and their change compared to the full list of m^6^A enriched transcripts (full methylome). We found that groups of transcripts were differentially represented in specific clusters (Fig. 4B). For example, RNA metabolism transcripts are more often methylated in stumpy forms only (*p*-value=0.02, two-sided Barnard’s unconditional test) (Fig. 4B, cluster 3). Interestingly, all detected methylated transcripts associated with nucleotide metabolism (N=5) are exclusively found in stumpy forms. In both slender and stumpy forms, VSG/VSG-related transcripts are found to be more often methylated than in procyclic forms, as well as genes associated with cellular processes (cluster 5) (*p*-value=0.01 and <0.01 respectively, two-sided Barnard’s unconditional test). Finally, transcripts associated with protein degradation and flagellum are more often methylated in the core methylome (cluster 1), than in any other cluster (*p*-value=0.02, two-sided Barnard’s unconditional test).

Overall, functional analysis of methylated transcripts shows that m^6^A methylation is found in transcripts encoding for a large variety of functions in the three stages of the parasite life cycle, and that each life cycle stage has a distinct functional methylome. Transcripts of a given functional class show multiple patterns of m^6^A methylation across the three life cycle stages, suggesting that m^6^A regulation acts in mRNAs encoded by individual genes and not in functional gene groups.

### Dynamics of m^6^A methylation in surface proteins

VSG expression is tightly controlled during the *T. brucei* life cycle. Our previous study in Lister 427 strain revealed that VSG transcripts are particularly enriched in m^6^A [16]. To investigate if VSGs are also enriched in EATRO1125 strain, we performed gene set enrichment analysis (GSEA) of VSG in slender, stumpy, and procyclic form transcriptomes (Fig. 5A, left panel). We detected significant enrichment for VSG in stumpy form m^6^A-IP samples (normalized enrichment score (NES) = 1.33, FDR *q*-value = 0.03). However, despite high enrichment scores in slender (NES=1.23) and procyclic form m^6^A-IP samples (NES=1.13), they did not pass the generally accepted statistical significance threshold for GSEA (FDR *q*-value = 0.25). As a negative control, we also performed GSEA on rRNA genes because m^6^A is less abundant in ribosomal RNA than in messenger RNA [10]. We confirmed our expectations that rRNA transcripts are significantly enriched in the input samples compared to m^6^A-IP samples (Slender forms NES= −1.08, FDR *q*-value = 0.10; Stumpy forms NES= −1.39, FDR *q*-value = 0.18; Procyclic forms NES= −1.33, FDR *q*-value = 0.17) (Fig. 5A, right panel). Overall, this data indicates that in EATRO1125, VSG remain a prominent class of methylated transcripts.

**Fig. 5.**
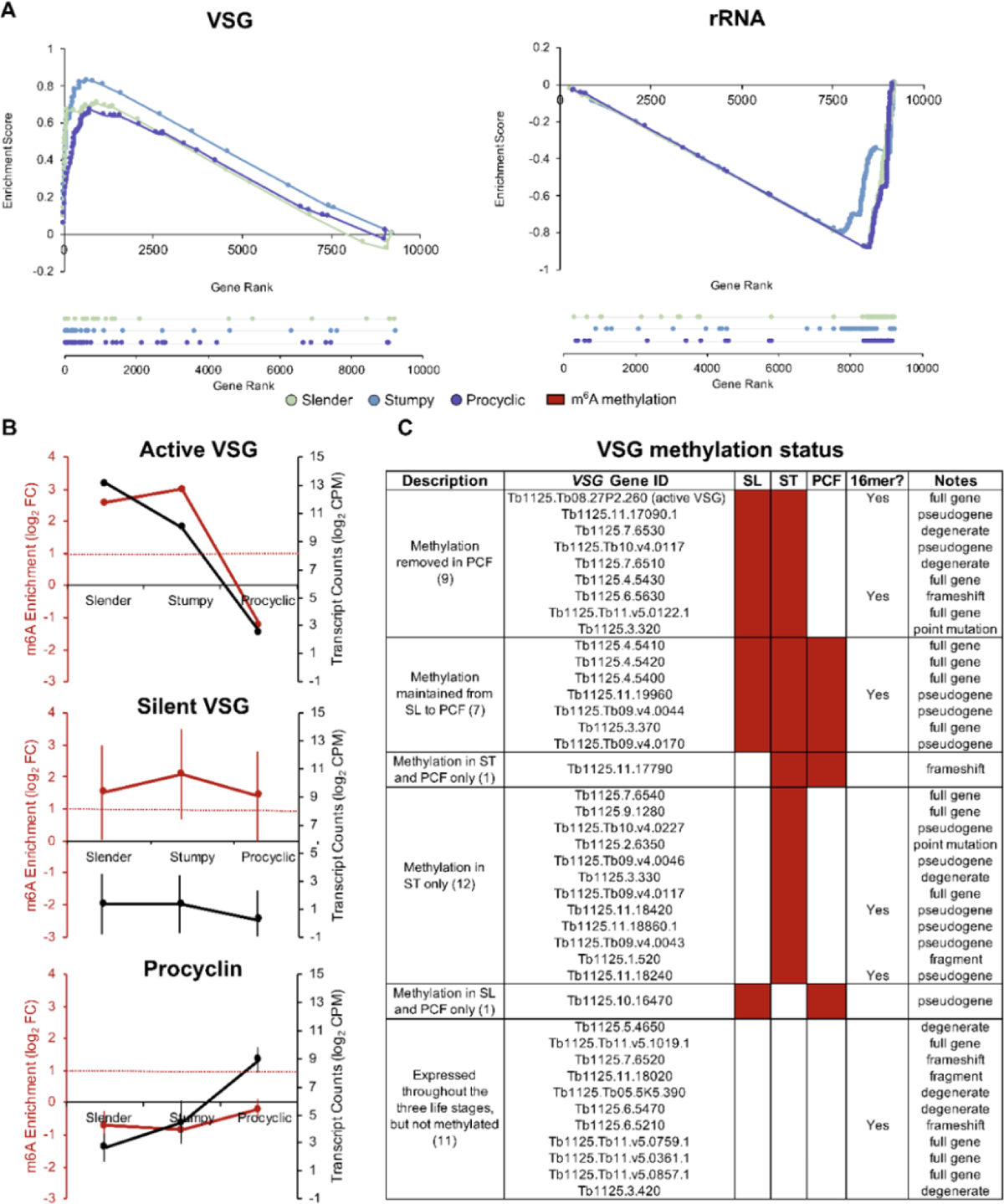
Variant surface glycoproteins (VSG) are enriched across *T. brucei* life cycle stages. **A**, Gene set enrichment analysis of VSG and rRNA in slender, stumpy and procyclic form parasites. **B**, m^6^A enrichment (red line, as log2 FC) and transcript counts (black line, as mean log-counts per million reads mapped) of the active VSG, silent VSGs and procyclins in slender, stumpy and procyclic form parasites. **D,** Schematic heatmap representing m^6^A methylation status of detected active and silent VSGs in slender (SL), stumpy (ST) and procyclic (PCF) forms and presence of 16-mer motif in their 3’UTR.

Next, we specifically inspected the behavior of the active VSG in the transition from slender, to stumpy, to procyclic forms. We confirmed that the expression of this transcript gradually decreased from highly abundant in slender forms to negligible in procyclic forms (Fig. 5B, first panel, black line). We further observed that the active VSG is methylated in slender and stumpy forms, but not methylated in procyclic forms (Fig. 5B, first panel, red line), suggesting a strong, positive correlation between m^6^A enrichment and transcript abundance (Pearson’s R^2^=0.86), and agreeing with our previous observations [16]. No other functional group or individual genes showed the same behavior, i.e. loss of methylation in procyclic forms associated to a sharp reduction in expression.

We then studied the methylation pattern of silent VSGs. For that, we compared the m^6^A enrichment levels in the 40 VSG transcripts detected in all the three stages of the life cycle. While silent VSGs have negligible expression levels throughout the life cycle (Fig. 5B, second panel, black line), their average enrichment in m^6^A-IP samples is high, suggesting that, unlike the active VSG, silent VSG transcripts may remain methylated in procyclic forms (Fig. 5B, second panel, red line). Next, we inspected the behavior of individual silent VSG transcripts. Of the 40 silent VSGs with detectable expression in all three life cycle stages, 16 were significantly methylated in slender forms, 28 in stumpy forms, and 9 in procyclic forms. Of those, 7 silent VSGs remain methylated throughout the three life cycle stages and 8 lose the methylation in procyclic forms. Interestingly, 12 VSGs are methylated exclusively in stumpy forms (Fig. 5C). We did not find a direct correlation between the presence of the conserved 16-mer motif in the 3’UTR of the detected VSGs and their methylation status. In summary, m^6^A enrichment in the silent VSG subset is variable and that m^6^A de-methylation is not always concomitant with differentiation to procyclic forms.

Finally, we investigated the behavior of EP procyclin transcripts (EP1-3 and GPEET) as they are hallmarks of procyclic form parasites [9, 34]. As expected, the transcript levels of EP procyclin was minor in slender and stumpy forms, but high in procyclic forms. In contrast to VSGs, we did not detect any enrichment for m^6^A methylation in EP procyclin transcripts, suggesting that they are not m^6^A methylated. These results show that not all major surface protein genes are regulated by m^6^A levels.

## Discussion

During a parasite’s life cycle, gene expression is tightly regulated to ensure parasites are best adapted to their environment. Post-transcriptional mechanisms are one way to regulate gene expression [2]. In this work, we hypothesized that m^6^A landscape varies across the life cycle as means of gene regulation. To test this, we compared the m^6^A landscape across three stages of the parasite’s life cycle: two proliferative forms (slender and procyclic forms) and one cell cycle-arrested form (stumpy forms). We found that methylation is more pervasive in stumpy forms, suggesting a role of methylation in the mechanisms that regulate quiescence.

In this study, we performed m^6^A immunoprecipitation without prior RNA fragmentation to obtain a complete set of methylated transcripts, regardless of the distribution of m^6^A modifications in poly(A) tails [16] or in internal locations [15]. Therefore, we did not assess which transcripts are internally or poly(A) methylated, and whether that impacts their expression and/or stability. However, in the future, mapping m^6^A location and stoichiometry within transcripts may be useful to clarify the role of this post-transcriptional modification in parasite development.

In mammalian cells, m^6^A methylation rarely changes across tissues or during different cell cycle stages [17, 18]. In contrast, we show that, in *T. brucei*, the core methylome represents only 21% of all detected transcripts. Each life cycle stage (slender, stumpy and procyclic forms) is characterized by a specific m^6^A methylome. We also found that the stumpy transcriptome is abundant in m^6^A-methylated transcripts (i.e 93% of all methylated transcripts are present in stumpy forms and 47% of methylated transcripts are exclusively found in stumpy forms), suggesting methylation of specific transcripts is important for the gene expression changes that characterize the stumpy forms. Given that in the growth-arrested stumpy form, transcription and translation are downregulated [7, 8], it is possible that m^6^A acts as a mechanism to stabilize critical mRNAs. It was previously shown in *T. brucei* that m^6^A in the poly(A) tail plays a stabilizing role in VSG transcripts and m^6^A methylation is associated with longer transcript half-life [15, 16]. Interestingly, several of the transcripts exclusively methylated in stumpy forms encode proteins involved in RNA metabolism (10%), which themselves could also contribute to mRNA stability in this short-lived life cycle stage. The reverse analysis of assessing the variation of the distribution of each functional group across each cluster, also reveals the same pattern, as transcripts involved in RNA metabolism fall 10% more within cluster 3 than any other cluster Finally, functional analysis of the core methylome itself shows that m^6^A is located in transcripts associated with different functions including flagellum and protein degradation, suggesting that m^6^A might affect various biological processes that are important in slender, stumpy and procyclic forms.

It was previously reported in Lister 427 strain that m^6^A methylation is important for VSG transcript stability [16]. Here, we confirmed that in an independent strain (EATRO1125) 41 VSG transcripts are also methylated. Interestingly, we found that in stumpy forms, two of the methylated transcripts encode for proteins critical for VSG expression control: VEX2 and CFB2. VEX2 (or VSG-exclusion protein 2) is part of the VEX complex, which is involved in VSG monoallelic expression by associating the VSG expression site with the spliced-leader array [35]. In *T. brucei* Lister 427, CFB2 stabilizes VSG transcripts by recognizing a conserved 16-mer motif in the VSG 3’ UTR and recruiting a protein complex that includes PABP2 [36]. Interestingly, the PABP2 transcript itself is also methylated and highly expressed throughout all life cycle stages. Human PABP was shown to increase translation in human cells [37]; in *T. brucei,* recent evidence indicates PABP2 might have a similar function for both VSG and bulk mRNA [36, 38]. Therefore, our results suggest a role for m^6^A in regulating VSG stability not only by methylation of VSG transcripts but also of transcripts that are important for VSG expression control in bloodstream forms.

In slender forms of Lister 427, transcript abundance of the active VSG was shown to be coupled to m^6^A methylation [16]. In this study, we confirmed this correlation in the three stages of the life cycle: in slender and stumpy forms the active VSG mRNA is abundant and m^6^A-methylated, while in procyclic forms the formerly active VSG mRNA is silenced and m^6^A levels drop. Demethylation of the active VSG transcript might be one m^6^A regulation mechanism triggered by differentiation, either by loss of inhibitory VSG complexes or by direct recruitment of demethylases.

Procyclin is the counterpart of VSG in procyclic forms, in terms of function (surface protein) and gene transcription (driven by RNA polymerase I) [39]. Researchers have therefore studied whether gene expression control of both VSG and procyclin extends beyond transcription or if at the post-transcriptional level, their regulation differs [40]. Here, we show that VSG is highly methylated while procyclin is not, suggesting distinct, stage specific, post-transcriptional mechanisms of gene regulation and highlighting the uniqueness of VSG expression control.

Silent VSGs are essential to maintain antigenic variation in slender forms by replacing the active VSG gene during VSG switching [41]. Surprisingly we found that among the silent VSGs, albeit negligible expression, 7 VSG genes are methylated in the three stages of the parasite life cycle, 8 VSG genes are methylated in slender and stumpy forms, but not in procyclic forms and 12 VSG genes are exclusively methylated in stumpy forms. These results point towards a differential regulation of m^6^A methylation dependent not only on the active or silent state, but also other factors that remain unknown.

Could m^6^A act as an epigenetic memory of VSG expression in *T. brucei*? In *Plasmodium falciparum*, monoallelic expression and switching of antigenic *var* genes are crucial for immune evasion of the parasite. Transmission of the active *var* gene from mature trophozoites to the next intraerythrocytic stage depends on a histone mark, H3K4me2, which is enriched at the promoter and allows for later transcription activation of the poised *var* gene [42, 43]. In *T. brucei*, multiple VSGs are transcribed by metacyclic form parasites in the tsetse fly before a single *VSG* gene is selected [44].

If methylation of VSGs associated with VSG switching and establishment of monoallelic expression, m^6^A could serve as a “memory mark” such that silent VSGs that remain methylated during differentiation from slender to procyclic forms would influence the choice of expressed VSGs at the metacyclic stage. Differentiation to epimastigote and metacyclic forms happens through a multi-stage process in the tsetse fly that includes re-activation of VSG expression and prepares the parasite for transmission to a mammalian host [45]. In the future, it will be interesting to identify which VSGs remain methylated in epimastigotes and metacyclic form parasites to test whether selective (de-)methylation contributes to the maintenance of different silent VSGs, preparing the parasite for future transmission.

In this work we provide the m^6^A landscape of *T. brucei* parasites from slender to procyclic forms, including transcripts methylated both internally and/or in the poly(A) tail. We show that the m^6^A landscape is life-cycle stage specific, contrasting with m^6^A regulation in mammalian cells that is more stable. We identified 491 stumpy-specific methylated transcripts that might be important to promote stumpy cell maintenance. Finally, we show that the dynamics of m^6^A methylation are different for surface proteins, suggesting differing regulation mechanisms for VSG and procyclin and within the VSG repertoire, which might play a role in VSG expression and/or selection across the parasite life cycle.

## Acknowledgments

The authors thank Leonor Pinho for technical and logistical assistance, the Genomics unit at Instituto Gulbenkian da Ciência for sequencing services and all members of the Figueiredo laboratory for helpful discussions and reagents. The project leading to these results has received funding from “la Caixa” Foundation under the agreement LCF/PR/HR20/52400019”. Researchers were funded by individual fellowships from FCT (2020.06827.BD to L.S.), MSCA ITN Cell2Cell (to L.LE). SSP received funding from Fundação Bial and Ordem dos Médicos through Prémio Maria de Sousa (7/2021) and FCT (2022.02187.PTDC), as well as the support of a fellowship from “la Caixa” Foundation (ID 10001043). I.J.V. was supported by “la Caixa” Foundation (HR20-00361) and H.M. by the European Research Council (ERC) under the European Union’s Horizon 2020 research and innovation programme (grant agreement no. 771714).

## Appendix A. Supplementary Data

Supplementary File A - Differential expression analysis including statistics for ST-MC vs ST-CPT (table A1); SL-MC (table A2); ST-MC (table A3); and PCF (table A4) Supplementary File B - m^6^A-Enriched and Depleted genes only in SL-MC (table B1); ST-MC (table B2); and PCF (table B3); m^6^A-Enriched genes in common with Viegas *et al.* dataset in 427 (table B4) Supplementary File C - Functional clustering of full methylome

